# Uncertain, intermittent access to reward promotes increased reward pursuit

**DOI:** 10.1101/2023.05.19.541519

**Authors:** Mike J.F. Robinson, Qi Shan A. Bonmariage, Anne-Noël Samaha

## Abstract

Self-administration procedures have been developed to model the intermittency of cocaine use in humans. These procedures involve intermittent, predictable access to cocaine during daily self-administration sessions. However, human drug use often involves intermittent and unpredictable patterns of drug access. Here, we introduce a new procedure to study the effects of unpredictable, intermittent access (UIntA) to a reinforcer, and we compare this procedure to two existing ones that provide predictable reinforcer availability; continuous (ContA) or intermittent (IntA) access. Three groups of rats self-administered water or a 5% sucrose solution in daily hour-long sessions. UIntA rats had alternating periods of reinforcer ON and OFF of unpredictable duration (1, 5 or 9 min/period). During reinforcer ON periods, reinforcer quantities were also unpredictable (0, 0.1 or 0.2 ml of solution) and were available under a variable ratio 3 schedule of reinforcement (1-6 responses). Both IntA and ContA rats had access to a fixed volume of water or sucrose (0.1 ml), under a fixed ratio 3 schedule of reinforcement. IntA rats had alternating and predictable 5-min reinforcer ON and OFF periods, while ContA rats had 60 minutes of reinforcer access during each session. Following 14 daily self-administration sessions, we found that UIntA rats had the highest rates of responding for water or sucrose reward under progressive ratio and extinction conditions, and the highest levels of cue-induced reinstatement of sucrose seeking. Thus, uncertain, intermittent access to reward promotes increased reward-seeking and -taking behaviours. This has implications for modeling addiction and other disorders of increased reward seeking.

## Highlights

- Human patterns of drug use involve uncertain and intermittent drug access
- We characterized a new model of uncertain intermittent reward access
- Uncertain intermittent access enhanced instrumental pursuit of sucrose and water
- This has implications for modeling disorders involving increased reward seeking

## 1. Introduction

A growing number of studies use intermittent access (IntA) self-administration procedures, to model patterns of drug seeking and taking thought to be relevant to addiction [1–4]. IntA involves intermittent drug availability, producing a cycle of peaks and troughs in brain concentrations of the drug [2,5–8]. IntA procedures not only more closely model human patterns of cocaine use, which are thought to be intermittent, both between and within periods of use [1,9–11], IntA is also uniquely effective in producing addiction-relevant features [1,3,4]. This includes persistent psychomotor sensitization [6,8,12], long-lasting enhancements in the motivation to obtain drug [5,6,13–16], and robust cue-induced drug seeking [14,16–20]. Though this work has often used cocaine, IntA experience with other drugs, including nicotine [21] and opioids [22–24], also promotes addiction-relevant patterns of drug taking and seeking.

Unlike human patterns of use which are often unpredictable and guided by several varying factors (drug availability, drug potency/purity, financial resources, etc.), IntA self-administration protocols provide very predictable patterns of reward access (typically 5 min ON and 25 min OFF, over 4-6-h sessions). Recent studies suggest that reward uncertainty can sensitize reward pathways, enhance motivation for reward and increase the ability of cues to trigger approach behavior [25–27]. For example, variable schedules of sucrose reinforcement produce greater dopamine release, locomotor cross-sensitization to amphetamine, and greater amphetamine self-administration, while uncertain reward outcomes promote increased attribution of incentive value to associated cues, recruit more environmental cues, and promote sign-tracking [28–31]. Reward uncertainty is also a crucial component of gambling and games of chance, and its role in rewarded behaviors highlights the cross-over between gambling and drug addiction [32–36].

Here we examined whether combining reward uncertainty and intermittent access to reward promotes greater reward seeking compared to predictable reward, as provided under IntA or continuous access (ContA) conditions. We hypothesized that uncertain intermittent access (UIntA) produces higher levels of responding for reward compared to IntA or ContA, and that IntA also enhances reward seeking and taking behaviours compared to ContA. To address these predictions, we allowed rats to self-administer water or a sucrose solution paired with discrete conditioned stimuli, under UIntA, IntA or ContA conditions. Following 14 daily self-administration sessions, we assessed responding for water or sucrose under a progressive ratio schedule of reinforcement and extinction conditions, as well as cue-induced reinstatement of water/sucrose-seeking behaviour.

## 2. Materials & Methods

### 2.1. Animals and Housing Conditions

Female Sprague-Dawley rats (N = 48; 125-150 g) purchased from Charles-River Laboratories (Kingston, NY, Barrier K90) were housed in pairs on a reverse 12h/12h light/dark cycle (lights on at 8:30 pm) with free access to food but restricted water access. Following 72 h of acclimation to the animal colony and before any experimentation, the rats were handled by experimenters once a day, for at least 3 days. Water restriction began with 6 h/day of water access for 4 days, followed by 4 h/day for the next 3 days, and then 2 h/day until the end of the experiments. Water was always given at least 1 hour after the end of testing, and at the same time each day. All procedures involving rats were approved by the animal ethics committee at the Université de Montréal and adhered to Canadian Council on Animal Care guidelines.

### 2.2. Groups and Conditions

Rats were randomly assigned to self-administer water or sucrose. Before the start of self-administration training, the rats received magazine training sessions to learn to retrieve experimenter-delivered water or sucrose from the magazine. Rats then received operant conditioning sessions as described below. Following operant conditioning, the rats were then assigned to one of 6 groups (Table 1).

**Table 1:** Rats were assigned to one of three different conditions following the acquisition of operant responding for water or sucrose, resulting in a total of 6 groups (3 for water and 3 for sucrose).

| **Group Name** | **Session Duration** | **Access period** | **No Access period** | **Total Access Period** | **Ratio** | **Reward Probability** | **Reinforcer volume** |
| --- | --- | --- | --- | --- | --- | --- | --- |
| **ContA** | 1 hr | 1 hr | 0 | 1 hr | FR3 | 100% | 0.1 ml |
| **IntA** | 1 hr | 5 min | 5 min | 30 min | FR3 | 100% | 0.1 ml |
| **UIntA** | 1 hr | 1, 5, or 9 min | 1, 5, or 9 min | 30 min | RR3 (1-6) | 33% | 0, 0.1, or 0.2 ml |

### 2.3. Behavioural Apparatus

Training and testing took place in standard operant chambers (31.8 x 25.4 x 26.7 cm; Med Associates, VT, USA. Illustrated in Fig. 1) located in a testing room separate from the colony room. The operant chambers were placed in sound-attenuating boxes equipped with a fan that reduced external noise. Each chamber contained a tone generator (2900 Hz; 85 dB) located adjacent to the house light on the back wall. On the opposite wall, there were two retractable levers on either side of a magazine equipped with an infrared head entry detector. A liquid dispenser was calibrated to deliver 100-µL drops of water or 5% sucrose into the magazine. Chambers were connected to a PC running Med-PC IV.

**Figure 1.**
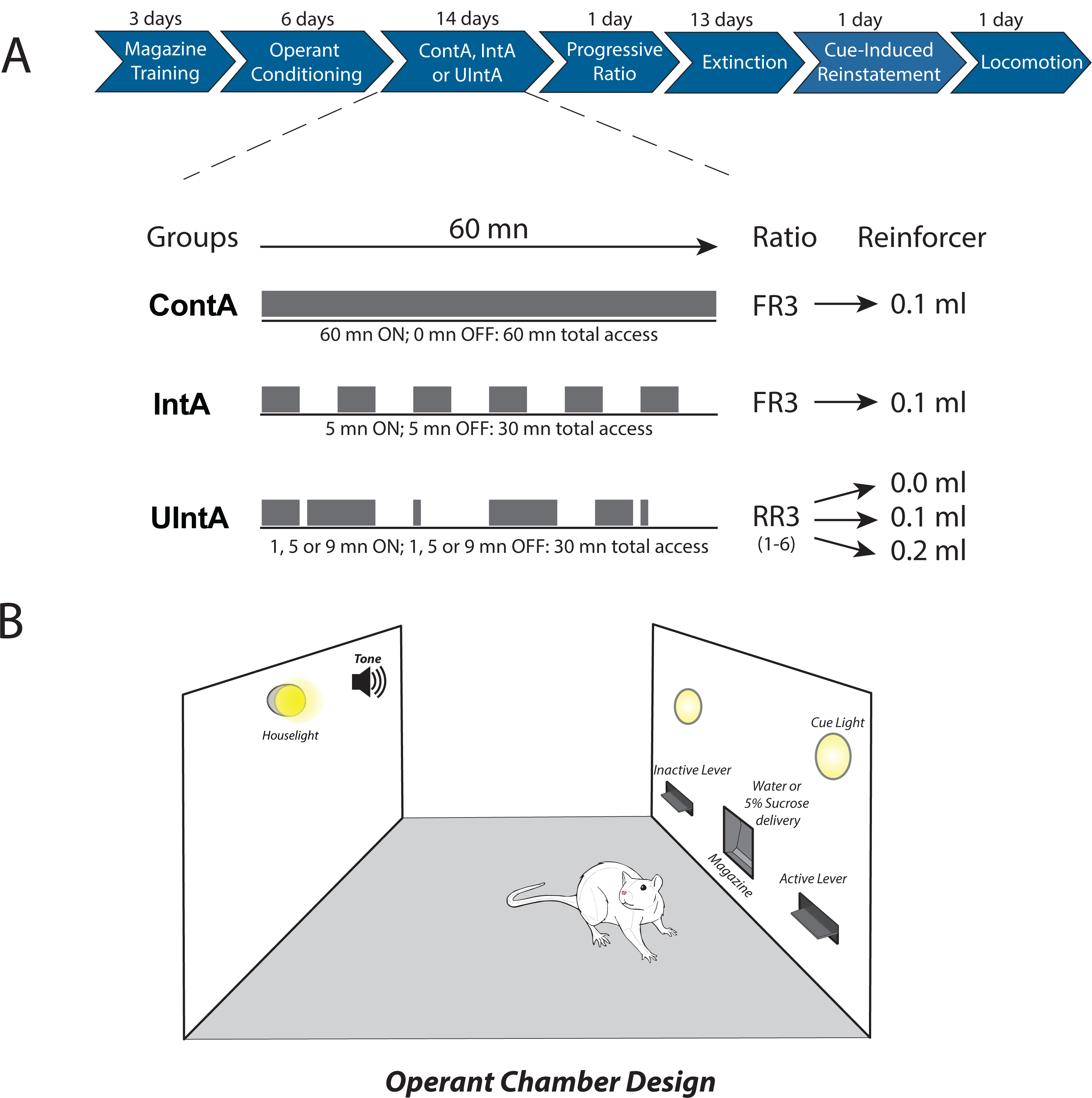
Experimental design and operant conditioning procedures. **A**, The sequence of training tasks and behavioural tests. After initial operant conditioning, rats were divided into 3 groups (Continuous Access – ContA; Intermittent Access – IntA; Uncertain Intermittent Access – UIntA) in each of 2 reinforcer conditions (water or 5% sucrose). The three groups differed on the pattern of reward access, the reinforcement ratio and reinforcer delivery conditions. **B**, The operant chambers included retractable active and inactive levers (counterbalanced), a magazine for liquid delivery, two cue lights above the levers and a tone generator and house light on the back wall of the chamber.

### 2.4. Procedures

#### 2.4.1. Magazine Training (3 sessions; 1 session/day)

Fig. 1A shows the sequence of experimental events. Two days prior to the start of magazine training sessions, rats in the Sucrose condition received 5% sucrose solution instead of water in their home cage during their daily, 2-h water access period, to reduce neophobia. Rats then received three consecutive daily magazine training sessions (30 min/session), where they were trained to retrieve experimenter-delivered water or sucrose from the magazine. Each rat was assigned to a specific operant chamber throughout all experimental procedures. The house light and fan were turned on to signal the beginning of each session. During each session, levers remained retracted and rats received thirty 100-µL deliveries of water or sucrose into the magazine under a variable interval 45 sec schedule (VI45s). Magazine head entries were recorded, and at the end of each session, the experimenters monitored whether any water/sucrose remained in the magazine.

#### 2.4.2. Operant conditioning (6 sessions; 1 session/day)

Following magazine training, rats received operant conditioning sessions during which they learned to press a lever for either water or sucrose (Fig. 1B). Rats were placed in the operant chambers with the two levers present. Responses on one (active) lever initially produced water or sucrose under a Fixed Ratio 1 schedule of reinforcement (FR1) for the first three days and then under FR3 for an additional three days. Responses on the other (inactive) lever were recorded but had no programmed consequence. The side of the active versus inactive lever was counterbalanced across rats. Levers always remained extended and there was no time out or cues presented following water or sucrose delivery. Sessions lasted 30 minutes or until an animal received 100 reinforcers, whichever came first. Rats needed to achieve a criterion of at least 2 days of a 2:1 ratio of active vs inactive lever presses to proceed from FR1 to FR3, and then to UIntA, IntA or ContA conditions.

#### 2.4.3. Sucrose/water self-administration under ContA, IntA, or UIntA conditions (14 sessions; 1 session/day)

At the end of operant conditioning, rats were assigned to the UIntA, IntA or ContA conditions (Fig. 1A), such that the sum of active lever presses made on the last 3 days of operant conditioning was comparable between groups. All self-administration sessions lasted 1 h and there was no limit on the number of reinforcers rats could earn per session. **ContA** rats had access to water/sucrose for the entire hour. **IntA** rats had access in 5-min periods (ON), followed by 5-min periods of no access (OFF), during which both levers were retracted. Thus, each 1-h session consisted of 6 cycles of intermittent ON/OFF access. For both ContA and IntA rats, water/sucrose was available under FR3, and each ratio completion was followed by a 6-s cue, consisting of illumination of the cue light directly above the active lever (either left or right) and a continuous tone (2900 Hz; 85 dB). During each 6-s cue presentation, levers remained extended, and responses on either lever were recorded but had no scheduled consequence (i.e., a timeout period). This conditioned stimulus (CS; 6 sec cue light+tone) was used in later tests for cue-induced reinstatement of extinguished reward-seeking behaviour. Finally, to achieve reward uncertainty in **UIntA** rats, during each session we introduced variability in *i*) the response ratio required, *ii*) the amount of water or sucrose delivered upon ratio completion, and *iii*) the length of each ON vs. OFF period. Thus, these rats had access to water/sucrose under a Random Ratio 3 schedule of reinforcement (RR3; 1-6 responses). Each ratio completion produced either 0, 100, or 200 µl of water/sucrose, each with a 33% probability. This meant that rats received on average 100 µl of water/sucrose, every 3 active lever presses. The light+tone CS (and 6-s timeout) was presented each time rats completed the FR3 requirement, whether water/sucrose was delivered or not. ON and OFF periods each lasted 1, 5, or 9 min (2 of each duration/period type, randomized). This produced a total of 6 cycles during each session, with an average of 5 min ON and 5 min OFF per cycle (see Fig 1A).

#### 2.4.4. Progressive ratio

Following operant conditioning, we measured responding for water/sucrose under a progressive ratio schedule of reinforcement, where the number of lever presses required for each successive water/sucrose delivery of the session increased according to the formula [5 × e (number of reinforcers×0.2) − 5] [37]. The active and inactive levers remained in the chambers during the session, and each ratio completion produced 100 µl water/sucrose, accompanied by the light+tone cue and 6-sec timeout. The session lasted an hour and breakpoint was calculated as the highest ratio completed within the session.

#### 2.4.5. Extinction (13 sessions)

After progressive ratio testing, rats received 1-h long extinction sessions (1 session/day) both to prepare rats for cue-induced reinstatement of lever pressing behaviour and to assess persistence in responding in the absence of reward. Here, both levers were presented throughout the session and lever pressing had no programmed consequences (no light+tone cue or water/sucrose delivery). Rats received daily extinction sessions until they reached an extinction criterion of <10% of the rate of active lever pressing seen on the last self-administration session.

#### 2.4.6. CS-induced reinstatement of extinguished water/sucrose-seeking

Immediately following the last extinction session, rats were tested for CS-induced reinstatement of water/sucrose-seeking behaviour. The reinstatement session lasted 1 hour and began with a response non-contingent presentation of the CS. Both levers were then inserted in the test cage and remained inserted throughout the session. Completion of the ratio (FR3) on the active lever resulted in CS presentation, but no water/sucrose delivery.

#### 2.4.7. Locomotor cross-sensitization to d-amphetamine

Exposure to reward uncertainty in the form of a variable ratio, as opposed to a fixed ratio, can produce locomotor cross-sensitization to amphetamine and increase dopamine release in the nucleus accumbens [31,38]. As such, following the test for CS-induced reinstatement of water/sucrose seeking, we tested our rats for locomotor cross-sensitization to d-amphetamine. Rats were placed in Plexiglas boxes (27 × 48 × 20 cm) equipped with 6 rows of photocells (3 cm above the box floor) to record spontaneous locomotion. Photocell counts were used as a measure of horizontal activity. After 30 minutes, rats received a saline injection (1 ml/kg, subcutaneously) and locomotion was recorded for a further 30 minutes. Finally, rats received d-amphetamine (1 mg/kg, subcutaneously; d-amphetamine sulfate, Sigma UK, via Health Canada; dissolved in 1 mg/ml saline) and locomotion was recorded for 1 more hour. The total number of photocell beam breaks was binned in 1-minute increments across the full 2-h session.

### 2.5. Statistical Analysis

Data from all tasks were analyzed using one-way or repeated measures ANOVAs or paired *t*-tests using SPSS 27 (IBM, Armonk, NY, USA), where appropriate. When interaction or main effects were statistically significant, post-hoc analyses (Fisher’s LSD) were conducted. Data are expressed as mean ± SEM. The *α* level was set at < 0.05. Where *p* values were < 0.001, we indicate ‘*p* = .000’.

## 3. Results

### 3.1. Magazine Training and Operant Conditioning

Prior to the start of operant conditioning, rats were initially given three days of magazine training to accustom them to the operant chambers and to retrieving water or sucrose) from the liquid delivery cup. Across reinforcer types, rats increased their number of magazine head entries across the three days (Fig 2A; Day: F_(2,68)_ = 10.22, *p* = .000). Compared to rats receiving water, rats receiving sucrose entered the magazine significantly more across days (Reinforcer: F_(1,34)_ = 17.43, *p* = .000; Reinforcer x Day: F_(2,68)_ = .56, *p* = .58). Thus, rats learned to retrieve water or sucrose in the magazine, and rats receiving sucrose visited the magazine more frequently.

**Figure 2.**
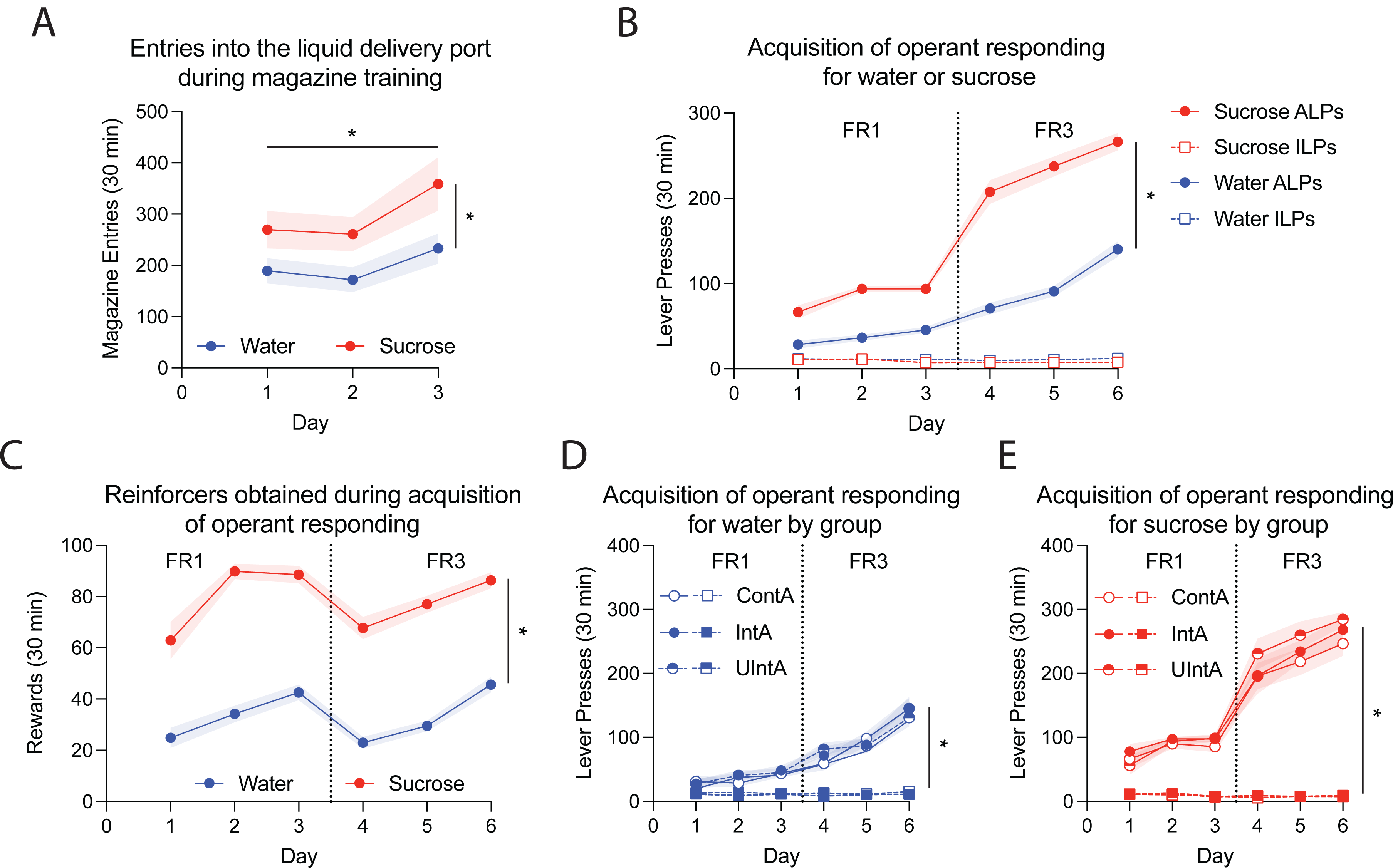
Rats acquire reliable water and sucrose self-administration behaviour, and respond more for sucrose than they do for water. **A**, Magazine entries during 30-min sessions during which rats received 30 experimenter-initiated water or sucrose deliveries (100-µL; VI45 s) across three consecutive days. **B**, Active (ALP) and Inactive (ILP) lever presses for water or sucrose across 3 days of fixed ratio 1 (FR1), followed by 3 days of FR3. **C**, The number of reinforcers obtained during each 30-min acquisition session. **D-E**, Lever presses during acquisition for water vs sucrose re-analyzed by respective group assignment. UIntA; Uncertain Intermittent Access. IntA; Intermittent Access. ContA; Continuous Access. **A-C**, n = 24/reinforcer. **D-E**, n = 8/group. * *p* < 0.05, main effect of Lever Type. Data are average +/-SEM.

Following magazine training, rats received operant conditioning sessions where they learned to lever press for water or sucrose, first under FR1 (Days 1-3), and then FR3 (Days 4-6). Across training days, rats press significantly more on the active versus inactive lever (Fig 2B; Day: F_(5,210)_ = 181.55, *p* = .000; Lever: F_(1,42)_ = 825.9, *p* = .000; Lever x Day: F_(1,42)_ = 215.71, *p* = .000). Compared to rats receiving water, rats receiving sucrose pressed more on the active lever, and showed a greater increase in active lever presses across days (Lever x Day x Reinforcer: F_(5,210)_ = 33.56, *p* = .000; Reinforcer: F_(1,42)_ = 121.47, *p* = .000; Day x Reinforcer: F_(5,210)_ = 27.18, *p* = .000; Lever x Reinforcer: F_(1,42)_ = 167.56, *p* = .000). Rats earned an increasing number of reinforcers across days (Fig 2C; Day: F_(5,210)_ = 19.5, *p* = .000), and they self-administered 2.5 times more sucrose than water (Reinforcer: F_(1,42)_ = 165.86, *p* = .000). Following operant conditioning, rats in each reinforcer condition were assigned to the UIntA, IntA or ContA groups such that the total number of active lever responses made on the last 3 instrumental conditioning days was comparable across groups. Retrospective analysis of acquisition data confirmed no group difference in responding for water or sucrose prior to group assignment (Fig 2D-E; Water: Group: F_(2,21)_ = .02, *p* = .98; Sucrose: Group: F_(2,21)_ = .79, *p* = .47). Thus, rats acquired water/sucrose self-administration behaviour, and rats given access to sucrose self-administered more. Moreover, under each reinforcer condition (water vs. sucrose), the 3 experimental groups had similar operant conditioning histories.

### 3.2. Sucrose/water self-administration under UIntA, IntA or ContA

After instrumental conditioning, rats continued to self-administer water or sucrose in daily sessions, but now under UIntA, IntA or ContA conditions. Both sucrose and water rats preferentially responded on the active lever, mostly ignoring the inactive lever (Fig. 3A; Lever: F_(1,21)_ = 354.20, *p* = .000; Fig. 3B; Lever: F_(1,21)_ = 149.90, *p* = .000). For each reinforcer type, there were no group differences in either lever-pressing behaviour (Water, Fig. 3A; Lever x Group: F_(2,21)_ = 1.95, *p* = .17; Group: F_(2,21)_ = 1.84, *p* = .18; Sucrose, Fig. 3B; Lever x Group: F_(2,21)_ = 1.97, *p* = .17; Group: F_(2,21)_ = 2.22, *p* = .13), or reinforcers earned (Water, Fig. 3A; Group: F_(2,21)_ = 0.19, *p* = .83; Sucrose, Fig. 3B; Group: F_(2,21)_ = 0.03, *p* = .97). Overall responding and preference for the active lever changed across days (Fig. 3A; Day: F_(13,273)_ = 4.27, *p* = .000; Lever x Day: F_(13,273)_ = 4.30, *p* = .000; Fig. 3B; Day: F_(13,273)_ = 13.05, *p* = .000; Lever x Day: F_(13,273)_ = 13.24, *p* = .000), and consequently, so did the number of self-administered reinforcers (Water, Fig 3C; Day: F_(13,273)_ = 3.74, *p* = .000; Sucrose, Fig. 3D; Day: F_(13,273)_ = 11.56, *p* = .000). Further analysis of this effect showed that rats trained with sucrose escalated their intake across days, with a primarily linear increase in consumption (F_(1,23)_ = 32.422, *p* = .000). In contrast, water trained rats showed primarily an Order 4 progression across days (F_(1,23)_ = 22.93, *p* = .000), suggesting variability in water self-administration across the two weeks of self-administration.

**Figure 3.**
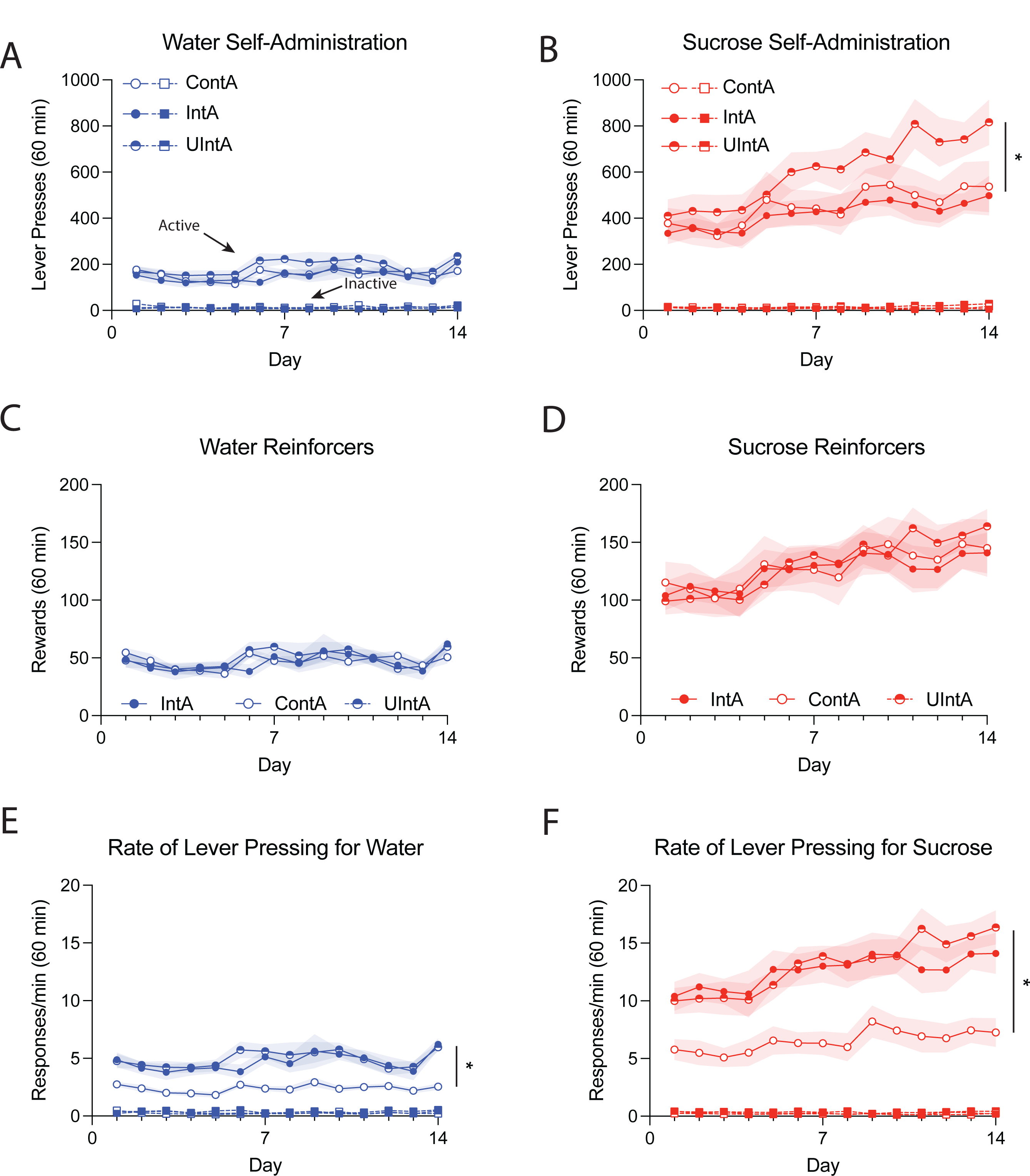
Access condition influences water and sucrose self-administration behaviour. **A-B**, Lever presses across 14 daily, 1-hour sessions of water or sucrose self-administration for Uncertain Intermittent (UIntA), Intermittent (IntA) and Continuous (ContA) access groups. **C-D**, The number of reinforcers earned during each self-administration session. **E-F**, The rate of responding for water or sucrose in the 3 access groups. n = 8/group. * *p* < 0.05, UIntA vs. other groups in **B**; ContA vs. other groups in **E-F**. Data are average +/-SEM.

We also analyzed the rate of lever pressing, defined as lever pressing per minute during periods of reinforcer availability, across days. Rats with intermittent access to water/sucrose (UIntA and IntA) responded at twice the rate of rats with continuous (ContA) access (Fig. 3E; Group: F_(2,21)_ = 15.34, *p* = .000: Fisher’s LSD: ContA vs IntA p = .000, ContA vs UIntA *p* = .000; Fig. 3F; F_(2,21)_ = 9.19, p = .001: Fisher’s LSD: ContA vs IntA *p* = .002, ContA vs UIntA *p* = .001). Thus, the intermittent groups responded at higher rates compared to the ContA groups, such that all groups earned similar numbers of reinforcers despite the intermittent groups having half the amount of time to respond per session compared to the ContA groups. This pattern of results was also consistent across reinforcer types, suggesting that intermittent access promotes increased rates of responding, regardless of the nature of the reinforcer (see also [39]).

Thus, both reinforcer type (water vs. sucrose) and access condition (UInta, IntA and ContA) influenced responding. Across access conditions, rats consumed more sucrose than water, and sucrose rats also escalated their intake across days. In parallel, access condition had no effect on the ability to discriminate between the active and inactive lever or on the amount of water/sucrose self-administered. However, access condition determined the rate of instrumental responding. Rats trained under intermittent access (UIntA and IntA) responded at twice the rate of ContA rats, allowing them to consume the same amount of water/sucrose.

The different access conditions produced distinct patterns of self-administration behaviour. Figure 4 shows the pattern of responding per minute for each group, across the two reinforcer types (sucrose in red vs water in blue) on the last self-administration day (Day 14). Levers were retracted during the OFF periods in the intermittent groups, but rats still managed to activate the retracted levers, albeit at low rates. Across access conditions and for both reinforcer types, rats responded most at the beginning of the session and reduced responding across the hour (Water, Figs. 4A-C; Minute: F_(59,1239)_ = 26.92, *p* = .000; Sucrose, Figs. 4D-F; Minute: F_(59,1239)_ = 17.28, *p* = .000), suggesting that rats experienced satiation. UIntA, IntA and ContA also produced distinctive patterns of lever pressing across the session (Water, Figs. 4A-C; Minute x Group: F_(118,1239)_ = 5.02, *p* = .000; Sucrose, Figs. 4D-F; Minute x Group: F_(118,1239)_ = 10.54, *p* = .000). UIntA produced an ON/OFF response pattern which was most apparent when the various ON/OFF periods (1, 5, 9 min) were aligned between rats, as done in Figs. 4A & D. IntA produced an ON/OFF response pattern which proceeded in predictable and orderly sequence (Figs. 4B & E). The IntA rats self-administering water also tended to load up on water at the beginning of each ON period and show a small but steady decline towards the transition to the OFF period (Fig. 4B; Comparing the 5 minutes of each ON period, Minute: F_(4,28)_ = 9.58, *p* = .000). In contrast, the IntA rats self-administering sucrose responded at comparable levels across each 5-min ON period (Fig. 4E; Minute: F_(4,28)_ = .68, *p* = .61). Finally, compared to the two other conditions, ContA produced a more sustained pattern of responding, until responding abated across the session (Figs. 4C & F).

**Figure 4.**
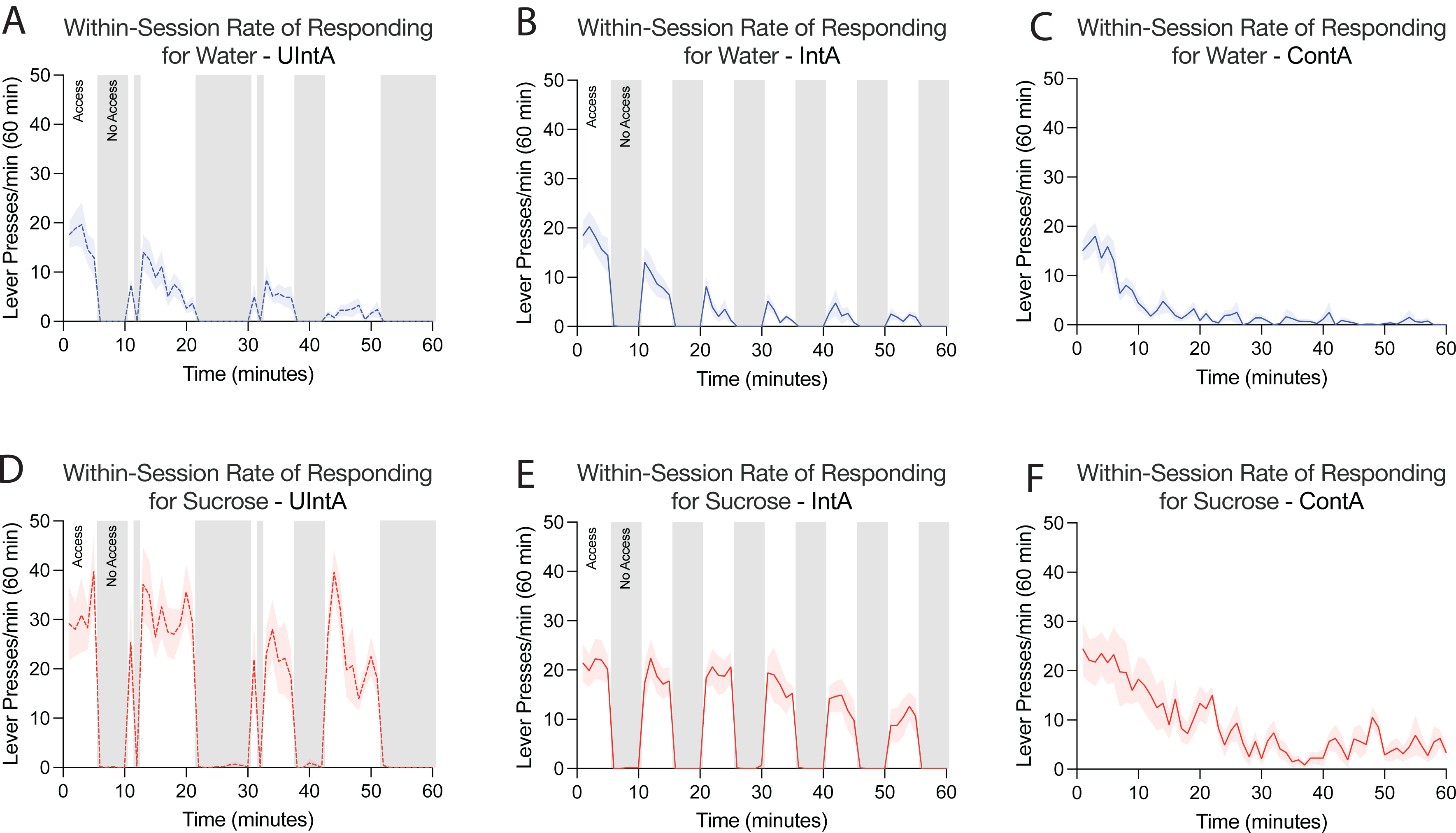
Access condition produces distinct self-administration profiles. **A**, Uncertain Intermittent access (UIntA) to water with the different periods of reinforcer access aligned across rats. **B**, Intermittent Access (IntA) to water. **C**, Continuous Access (ContA) to water. **D**, UIntA to sucrose with the different periods of reinforcer access aligned across rats. **E**, IntA to sucrose. **F**, ContA to sucrose. Gray shading indicates periods of reinforcer unavailability. n = 8/group. Data are average +/-SEM.

Figure 5 shows the rate of active lever pressing specifically during the 6-s presentation of the reward-associated CS. Although responding had no programmed consequence, it can be indicative of reward seeking during cue presentation, and an animal’s general degree of reward-seeking motivation. As each CS presentation lasted 6 s, across training sessions, the number of responses during CS presentation accounted for only a small fraction (Water approx. 10%; Sucrose approx. 16%, data not shown) of the total responses during each 1-h session. Notably, rats increased their responding on the active lever during CS presentation across sessions, even though responding continued to have no programmed consequence (Fig 5A; Day: F_(13,273)_ = 6.72, *p* = .000; Fig 5B; F_(13,273)_ = 7.81, *p* = .000).

**Figure 5.**
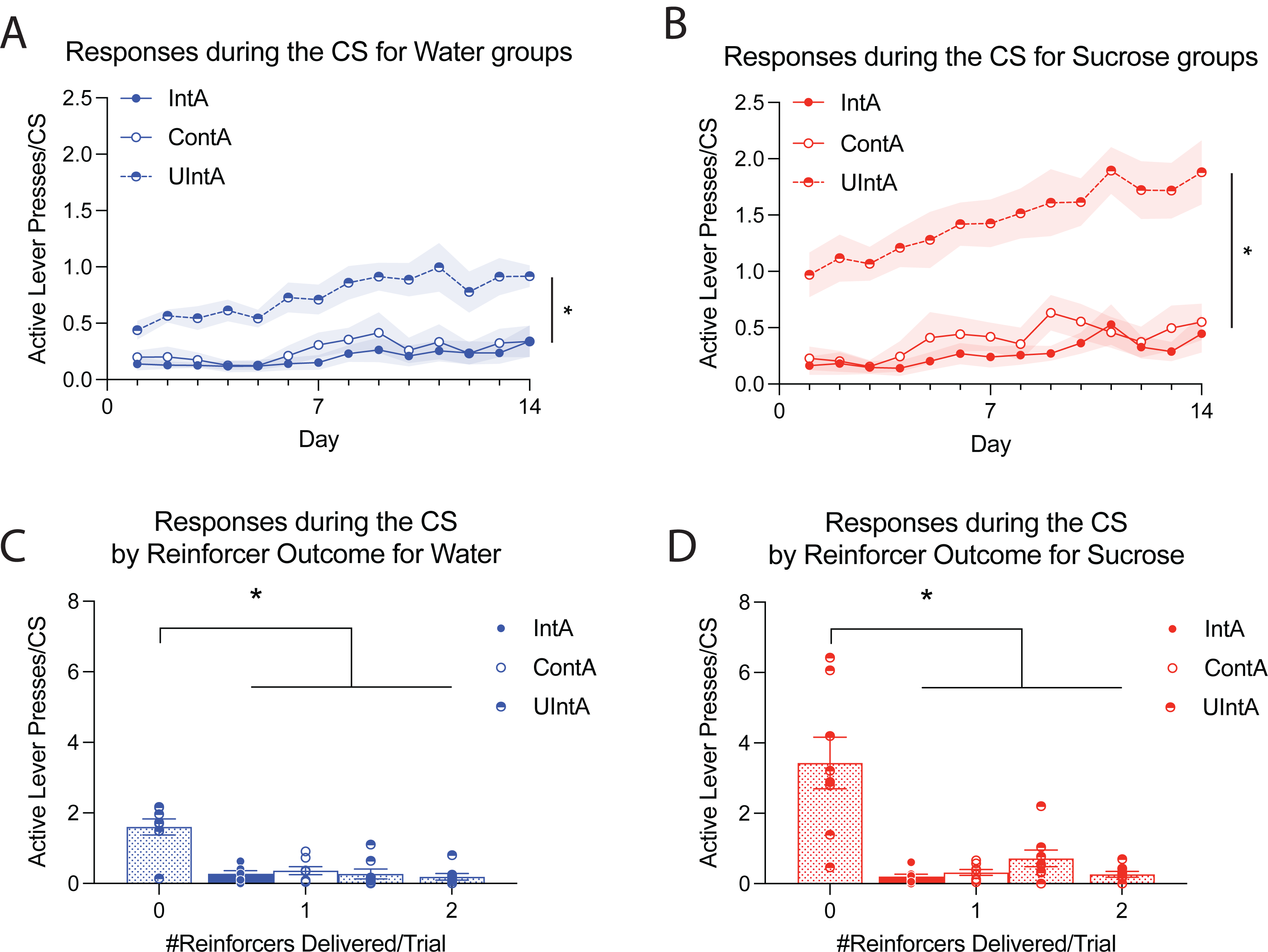
Rats given Uncertain Intermittent Access (UIntA) to water or sucrose show greater lever-pressing behaviour during conditioned stimulus (CS) presentation compared to the other groups. **A-B**, Active lever presses per second per 6-second CS presentation for water and sucrose across the 14 days of self-administration. **C-D**, For water and sucrose groups, respectively, total responses per each 6-second CS presentation when 0, 1 or 2 reinforcers were delivered (100 uL each delivery). As detailed in the text, in the IntA and ContA conditions, each CS presentation was always followed by delivery of 1 reinforcer, whereas in the UIntA condition, each CS presentation could be followed by 0, 1 or 2 reinforcers. IntA; Intermittent Access. ContA; Continuous Access. n = 8/group. * *p* < 0.05, UIntA vs. other groups in **A-B**. Data are average +/-SEM.

Lever-pressing behaviour during CS presentation depended on access condition. UIntA rats pressed significantly more on the active lever during CS presentations compared to the other groups (Fig. 5A - Water; Group: F_(2,21)_ = 12.43, *p* = .000; Fig. 5B - Sucrose; Group: F_(2,21)_ = 25.43, *p* = .000; Fisher’s LSD UIntA *p*’s ≤ .001). UIntA rats had an equal probability of receiving 0, 1 or 2 reinforcers each time they completed the response ratio (FR3). We sought to determine whether the number of reinforcers delivered on each ratio completion (or trial) influenced active lever presses during CS presentation (Figs. 5C-D). Indeed, it did. During trials when UIntA rats earned 1 or 2 reinforcers, UIntA rats performed a similar number of active lever presses during CS presentation as did the other groups (Water, Fig. 5C; F_(2,21)_ = .659, *p* = .528; Sucrose, Fig. 5D; F_(2,21)_ = 3.211, *p* = .061). However, during unreinforced trials, UIntA rats significantly increased the rate of these CS responses, both compared to trials when UIntA rats earned 1 or 2 reinforcers and compared to the other groups (Water, Fig. 5C; #Reinforcers delivered/trial: F_(2,14)_ = 29.56, *p* = .000; 0 reinforcer vs 1 or 2 reinforcers: *t*_(7)_’s > 5.66, all *P*’s < .001; Sucrose, Fig. 5D; #Reinforcers delivered/trial: F_(2,14)_ = 14.26, *p* = .000; 0 reinforcer vs 1 or 2 reinforcers: *t*_(7)_’s >3.35, all *P*’s < .007). Thus, reward uncertainty, and in particular the occasional absence of reward, promotes reward seeking in the presence of the reward-associated CS.

### 3.3. Progressive Ratio

Following the 14 UIntA, IntA or ContA sessions, we compared the groups on responding for water/sucrose under a progressive ratio schedule of reinforcement. Across groups, rats pressed significantly more on the active vs. inactive lever (Fig. 6A; Water: Lever: F_(1,21)_ = 93.16, *p* = .000; Fig. 6B; Sucrose: Lever: F_(1,21)_ = 196.11, *p* = .000). In addition, animals trained with sucrose pressed more on the active lever and reached higher breakpoints than those trained with water (Water vs Sucrose: Active Responses: F_(1,47)_ = 18.05, *p* = .000; Breakpoint: F_(1,47)_ = 17.41, *p* = .000). There were also group differences in responding. UIntA rats lever pressed more than IntA and ContA rats did, for both water (Fig. 6A; Group: F_(2,21)_ = 4.43, *p* = .025; Fisher’s LSD: UIntA vs IntA *p* = .013, UIntA vs ContA *p* = .03) and sucrose (Fig. 6B; Group: F_(2,21)_ = 21.43, *p* = .000; Fisher’s LSD: UIntA vs IntA *p* = .000, UIntA vs ContA *p* = .000). In addition, the UIntA rats also showed stronger discrimination between the active vs. inactive lever, at least in the sucrose (but not the water) condition (Fig. 6A; Lever x Group: F_(2,21)_ = 3.13, *p* = .065; Fig. 6B; Lever x Group: F_(2,21)_ = 20.07, *p* = .000).

**Figure 6.**
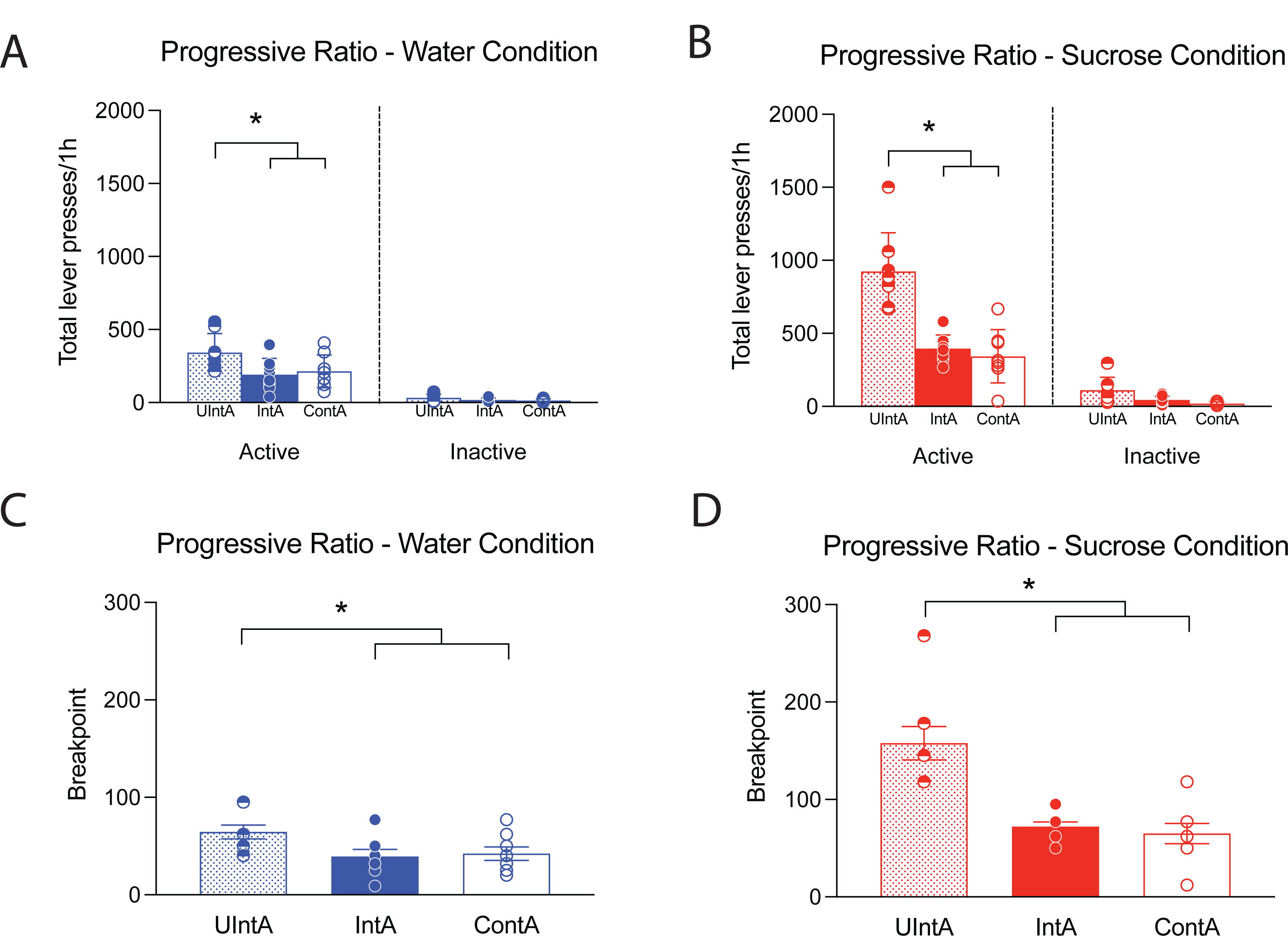
Rats given Uncertain Intermittent Access (UIntA) to water or sucrose later respond more for these reinforcers under progressive ratio conditions. **A-B**, Total number of presses on the active and inactive lever as a function of access condition and reinforcer type (water or sucrose) during a single 1-hour progressive ratio session. **C-D**, Breakpoint reached as a function of access condition. n = 8/group. * *p* < 0.05. Data are average +/-SEM.

Accordingly, UIntA rats achieved the highest breakpoints for both water (Fig. 6C; Group: F_(2,23)_ = 3.76, *p* = .04; Fisher’s LSD: UIntA vs IntA *p* = .021, UIntA vs ContA *p* = .04) and sucrose (Fig. 6D; Group: F_(2,23)_ = 18.77, *p* = .000; Fisher’s LSD: UIntA vs IntA *p* = .000, UIntA vs ContA *p* = .000). IntA and ContA did not differ from one another. Thus, under conditions where obtaining water or sucrose cost increasing physical effort, rats with a history of uncertain, intermittent access to these reinforcers (UIntA) worked harder to obtain them than rats with a history of predictable reinforcer access (i.e., IntA or ContA rats).

### 3.4. Extinction and Cue-Induced Reinstatement

After progressive ratio testing, rats received extinction sessions, during which responses on either lever had no programmed consequences. It took 13 sessions for all rats to meet the extinction criterion of active lever responding below 10% of what it was on the last day of water/sucrose self-administration (Day 14 in Figs. 3A-B). Across these 13 days of extinction, rats trained with either water or sucrose significantly decreased their lever-pressing behaviour (Fig. 7A; Day: F_(12,252)_ = 21, *p* = .000; Fig. 7B; Day: F_(12,252)_ = 26.97, *p* = .000), in particular on the active lever (Fig. 7A; Day x Lever: F_(12,252)_ = 27.17, *p* = .000; Fig. 7B; Day x Lever: F_(12,252)_ = 36.96, *p* = .000). There were also group differences in extinction responding. Under both water and sucrose conditions, UIntA rats showed a slower rate of extinction than IntA or ContA rats did (Fig. 7A; Group: F_(2,21)_ = 8.23, *p* = .002; Group x Day: F_(12,252)_ = 1.63, *p* = .04; Fisher’s LSD: UIntA vs IntA *p* = .004; UIntA vs ContA *p* = .001; IntA vs ContA *p* = .677; Fig. 7B; Group: F_(2,21)_ = 15.69, *p* = .000; Group x Day: F_(12,252)_ = 3.82, *p* = .000; Fisher’s LSD: UIntA vs IntA *p* = .000; UIntA vs ContA *p* = .000; IntA vs ContA *p* = .794). Thus, after a history of UIntA to reward, rats will persist for longer in seeking reward under conditions where it is no longer available.

**Figure 7.**
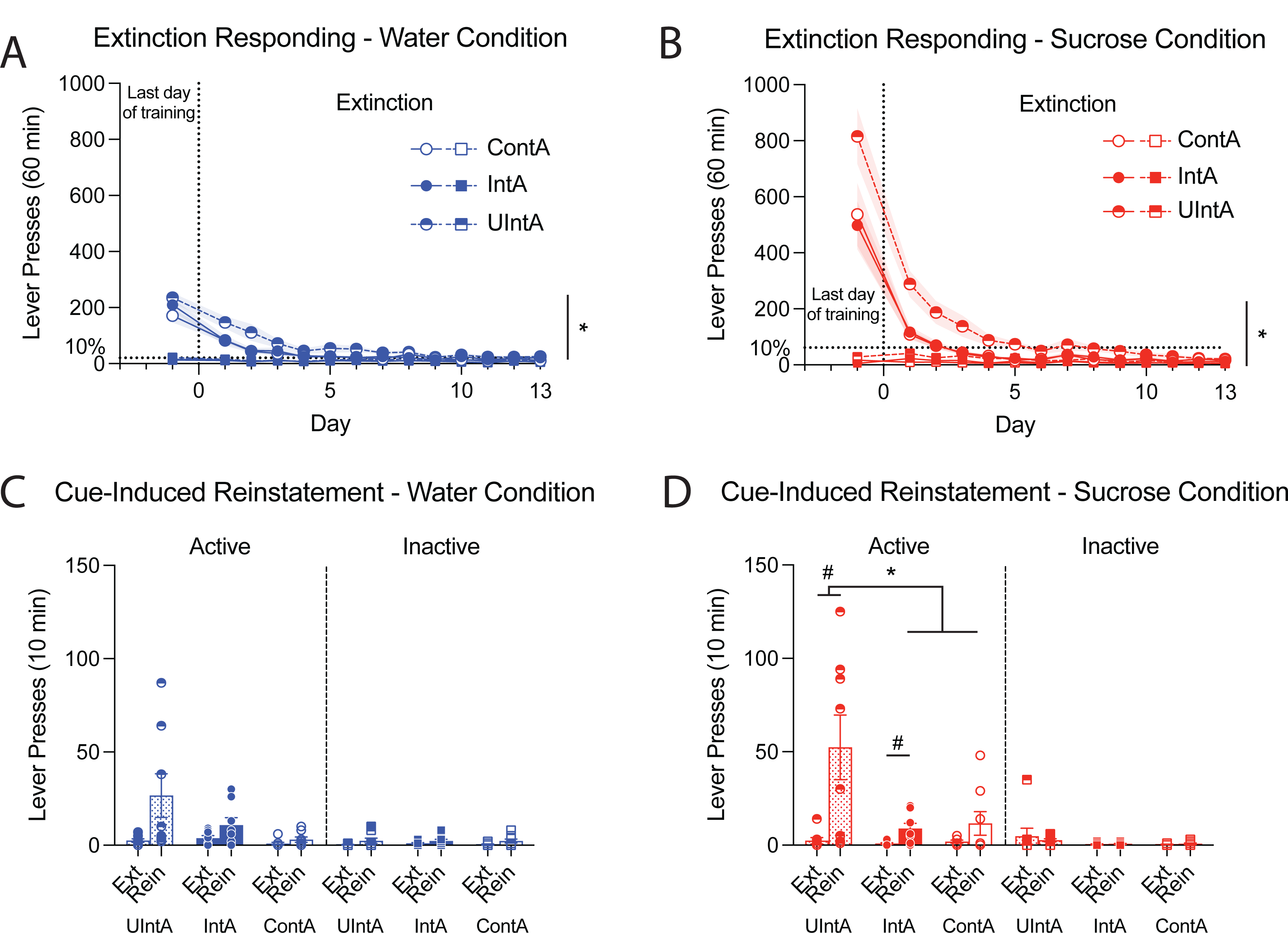
Rats given Uncertain Intermittent Access (UIntA) to water or sucrose later respond more under extinction conditions and also show more robust conditioned stimulus-induced reinstatement of reward-seeking behaviour compared to the other groups. **A-B**, Lever responding on the last day of access conditioning, compared to responding on 13 consecutive days of extinction training where responses on either lever had no programmed consequence (no reinforcer, no conditioned stimulus). **C-D**, Lever pressing during the last 10 minutes of the last extinction session (‘Ext.’) vs the first 10 minutes of the conditioned stimulus-induced reinstatement session (‘Rein’). n = 8/group. * *p* < 0.05, Active lever presses in the UIntA vs. other groups in **A-B**; Reinstatement responding in the UIntA vs. other groups in **C-D**. # *p* < 0.05, Extinction vs. Reinstatement within the same group in **D**. Data are average +/-SEM.

Immediately after the final extinction session, rats were tested for CS-induced reinstatement of reward-seeking behaviour (1-h session). During reinstatement tests, rats responded at high levels in the first 10 min and decreased responding thereafter (data not shown). Responding was therefore compared between the first 10 min of the reinstatement session and the last 10 min of the final extinction session. Collapsing across access conditions, rats pressed more on the active vs. inactive lever in both water and sucrose conditions (Fig. 7C; Water: Lever: F_(1,21)_ = 9.971, *p* = .005; Fig. 7D; Sucrose: Lever: F_(1,21)_ = 12.904, *p* = .002). Collapsing across access conditions, the rats also pressed more on the active lever during the reinstatement session compared to the last extinction session (Fig. 7C; Test x Lever: F_(1,21)_ = 6.002, *p* = .023; Fig. 7D; Test x Lever: F_(1,21)_ = 15.262, *p* = .000). In sucrose-trained rats, active lever presses varied as a function of test type (extinction vs. reinstatement), lever type and access group (Fig. 7D; Sucrose: Test x Lever x Group: F_(2,21)_ = 6.046, *p* = .008; a similar, nonsignificant trend emerged for water-trained rats, Fig. 7C; Test x Lever x Group: F_(2,21)_ = 2.937, *p* = .075). Further analysis of data from the sucrose-trained rats showed that only the intermittent access groups (IntA & UIntA) significantly increased active lever responding during the reinstatement session compared to the extinction session (Fig. 7D, Test x Lever; UIntA: F_(1,7)_ = 10.002, *p* = .016; IntA: F_(1,7)_ = 9.792, *p* = .017; For Active lever presses: Test: UIntA: F_(1,7)_ = 9.143, *p* = .019; IntA: F_(1,7)_ = 8.168, *p* = .024). Crucially, only the UIntA group showed significantly more CS-induced reinstatement compared to the other groups (450% & 590% active lever press increase over ContA and IntA, respectively. Test x Group: F_(2,21)_ = 4.832, *p* = .019; Fisher’s LSD: UIntA vs IntA *p* = .008, UIntA vs ContA *p* = .013). Thus, presentation of the CS reinstated extinguished reward-seeking behaviour in rats that have experienced intermittent reward access (UIntA and IntA groups), and did so significantly more in those exposed to uncertain (UIntA) rather than predictable reward access (IntA or ContA).

### 3.5. D-Amphetamine-induced Locomotion

Finally, we examined whether different histories of water/sucrose self-administration produced cross-sensitization to the psychomotor activating effects of d-amphetamine. Figs. 8A-B show the locomotor response to saline- and d-amphetamine. Initial locomotion was relatively high, likely due to exploration of the novel environment. Locomotor activity decreased thereafter, such that rats showed less locomotion in the 2^nd^ half-hour of the test vs. the first (Figs 8A-B; first 30 min vs Saline: Water: F_(1,21)_ = 64.79, *p* = .000; Sucrose: F_(1,21)_ = 26.47, *p* = .000). D-amphetamine increased locomotion relative to saline (Saline vs Amphetamine: Fig. 8A, Water: F_(1,21)_ = 199.23, *p* = .000; Fig. 8B, Sucrose: F_(1,21)_ = 228.07, *p* = .000), however, there was no effect of access condition on d-amphetamine-induced locomotion (Group: Water: all *P*’s > .22; Sucrose: all *P*’s > .13). Thus, under our conditions, uncertain reward access in the UIntA rats did not change the locomotor activating effects of d-amphetamine.

**Figure 8.**
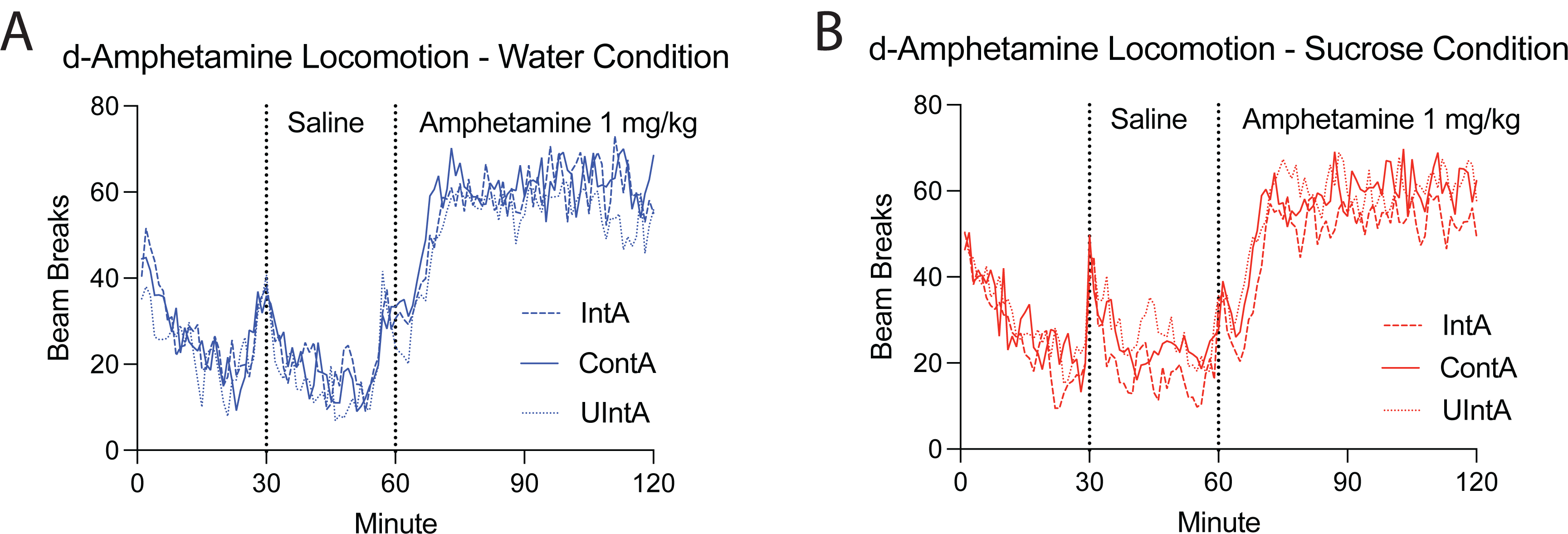
Locomotor activity under baseline conditions and in response to a saline then a D-amphetamine injection. **A-B**, Spontaneous locomotion (30 mins) measured in beam breaks for water vs sucrose rats, followed by a saline injection (30 mins) and an injection of d-amphetamine (60 mins).

## 4. Discussion

We examined the hypothesis that reward uncertainty during intermittent access (UIntA) enhances reward-seeking and -taking behaviours relative to predictable reward, delivered either on an intermittent or continuous schedule (respectively, IntA or ContA). To introduce uncertainty, we varied the duration of reinforcer access and non-access periods, the effort required to get reinforcement and the probability and magnitude of reinforcement. To test our hypothesis, we also compared two reinforcer types; water and liquid sucrose. We report four main findings. First, access condition did not influence the amount of water or sucrose rats self-administered. However, whether with reward certainty (IntA) or uncertainty (UIntA), intermittent access produced significantly higher rates of responding during the self-administration sessions. Thus, UIntA and IntA rats earned a similar number of reinforcers as ContA rats did, but in half the session time. Second, during the self-administration sessions, UIntA rats responded more during CS presentation than did the other groups, in particular on trials where CS presentation was not followed by reinforcer delivery. Third, compared to ContA or IntA rats, UIntA rats displayed increased rates of responding both under progressive ratio and extinction conditions, suggesting enhanced motivation for water or sucrose (also see [38,40]). Fourth, only intermittent access rats (IntA & UIntA) showed significant CS-induced reinstatement of extinguished sucrose seeking, and UIntA rats showed a 4x greater increase in reinstatement behaviour compared to IntA and ContA rats.

### 4.1. Effects of continuous vs. intermittent access

Intermittent access to water or liquid sucrose (UIntA and IntA) promoted higher rates of instrumental responding during self-administration sessions compared to ContA. This extends prior work using liquid saccharin [39], and indicates that when the active lever is present, rats with intermittent access to reinforcement experience higher rates of reinforcement as compared to ContA rats. However—and to our surprise— under classical IntA conditions, this higher rate of reinforcement did not translate to greater breakpoints achieved for water or sucrose during progressive ratio tests or greater responding under extinction conditions. Similarly, while within-subjects analyses showed significant CS-induced reinstatement of sucrose seeking behaviour in IntA rats, but not in ContA rats, there were no significant differences when the groups were compared directly. This is in contrast to studies showing that compared to ContA, IntA to cocaine [1,3,7,14,16,17,20], opioids [22,23,41] or nicotine [21] is especially effective in promoting the pursuit of drugs. Intermittent drug access produces repeated spikes in brain drug concentrations, and these are thought to trigger sensitization-related neuroadaptations that then cause increased motivation for drug [2–4,6,7,12,42]. Given this, our findings suggest that intermittent intake of non-drug rewards does not produce these neuroadaptations, because ContA vs. IntA produced similar levels of responding for water and sucrose during progressive ratio and extinction tests. Thus, the effects of intermittency on reward-seeking behaviour might not robustly generalize to non-drug reinforcers.

Our findings relate perhaps most directly to the work of Beasley and colleagues [39]. Their findings show that compared to continuous access, IntA to saccharin produces higher breakpoints during progressive ratio tests. Our findings are in apparent contradiction, as our ContA and IntA rats achieved similar breakpoints for water as well as for sucrose. Yet several procedural differences distinguish our two studies. First, Beasley et al used saccharin [39] and we used sucrose. However, we do not believe this was decisive. Their saccharin is calorie-free just like our water reinforcer, and their saccharin has a sweet taste just like our sucrose reinforcer. Importantly, our results were constant across reinforcer types, showing no differences between ContA and IntA groups on breakpoints or reward seeking under extinction. Second, their rats had 6-hour self-administration sessions, rather than the 1-hour sessions used here. Our shorter sessions could have blunted the effects of intermittency on later motivation for reward. This is possible. However, we have previously shown that longer (6 h) versus shorter (2 h) IntA sessions produce similar responding under both progressive ratio and extinction conditions, at least for cocaine [7]. Third, their 6-h IntA sessions involved 5-min reinforcer ON periods separated by 25-min reinforcer OFF periods. The longer OFF periods may have promoted greater levels of craving due to the more prolonged periods of reinforcer unavailability. This could have resulted in more motivation for saccharin when effort rather than time was the primary constraint on reinforcer availability during the progressive ratio task. Finally, the IntA condition provided a 1:1 ratio of access time in our study (5 min ON, 5 min OFF) vs. a 5:1 ratio (5 min ON, 25 min OFF) in Beasley et al [39]. In parallel, our IntA rats earned the same number of reinforcers as our ContA rats, presumably reaching the same degree of satiation. This could explain the lack of differences in later breakpoints, extinction responding and reinstatement between our IntA and ContA rats.

Beasley et al [39] propose that in addition to drug-related neuroadaptations, IntA promotes increased motivation for reward because under IntA conditions, the lever is a discriminative cue associated with a higher rate of reinforcement. This in turn could result in more Pavlovian excitation being conditioned to the lever after IntA versus ContA training, thus producing increased reward-seeking and -taking behaviours. Their results are quite consistent with this hypothesis, but ours are not. Here, IntA (and UIntA) rats showed higher rates of responding compared to the ContA group. Yet, IntA and ContA rats were similar on measures of reward seeking and taking. This argues against a Pavlovian excitation explanation of the effects of intermittency. It is also unlikely that greater Pavlovian excitation conditioned to the lever with IntA vs. ContA procedures can explain the other effects produced by IntA drug access, such as psychomotor and dopamine sensitization [6,12,15,16,43].

### 4.2. Effects of reward uncertainty under intermittent access conditions

Our findings support an exacerbating role of uncertainty on reward seeking (see also [38,40]). First, compared to IntA and ContA groups, UIntA rats achieved higher breakpoints for water and sucrose and greater resistance to extinction when reinforcement was no longer available. Several aspects of the UIntA procedure could explain these effects. First, during prior self-administration sessions, UIntA rats were under a variable ratio schedule of reinforcement, while IntA and ContA rats were under a fixed ratio schedule. This could have promoted continued responding in the UIntA rats when reinforcement is suddenly costly (i.e., during progressive ratio tests) or not available (during extinction). This is also in line with the increased persistence of responding effects that uncertainty has during omission training and extinction under Pavlovian conditions [44,45]. The UIntA rats also showed evidence of enhanced sensitivity to the ability of cues to motivate instrumental responding, akin to Pavlovian-to-instrumental transfer [46,47]. Indeed, compared to the other groups, our UIntA rats showed greater reward-seeking behaviour when the reward-associated CS was presented during the self-administration sessions (Fig. 5), even though this represented a timeout period, with no reinforcer available. UIntA rats also showed significantly more CS-induced reinstatement of extinguished reward seeking as compared to IntA or ContA rats. Prior studies suggest that reward uncertainty enhances incentive salience attribution and can even recruit otherwise ignored or less salient cues and assign them with incentive value, despite reduced predictive value [28,29,48,49]. Thus, our findings are consistent with the idea that reward uncertainty promotes the attribution of incentive value to reward-associated cues [25,49], thereby increasing the ability of these cues to trigger and invigorate reward seeking actions.

There were no effects of reward uncertainty (or intermittency) on d-amphetamine-induced locomotion. Extended exposure to an uncertain reward can sensitize brain reward pathways and thus promote cross-sensitization to the psychomotor activating effects of stimulant drugs [31,38]. However, the protocol used in previous studies involved a far greater degree of reward uncertainty than ours in terms of instrumental responding (from VR1 to VR20), and previous studies also involved a more extended training period (55 days vs 14 days here). Further work testing different parameters (more extended self-administration, additional reinforcers and abstinence periods) is needed to conclusively determine whether our UIntA protocol promotes cross-sensitization to the psychomotor activating effects of stimulant drugs.

### 4.3. Conclusion

We provide a new procedure to study the behavioural and neurobiological effects of uncertain, intermittent access to reward. This procedure arguably models naturalistic conditions, with ecologically valid levels of unpredictability, whereby reward availability is intermittent and unpredictable. Using this procedure, we demonstrate that—across reinforcer types— factors associated with reward uncertainty increase both motivation for reward as well as the ability of reward-associated cues to gain control over appetitive behaviour. This has implications for modeling several disorders characterized by pathological reward-seeking behaviour, including drug addiction, eating and gambling disorders.

## Acknowledgements

This work was supported by the National Science and Engineering Research Council of Canada (grant 355923) and the Canada Foundation for Innovation to ANS (grant 24326). ANS holds a salary award from the Fonds de la Recherche du Québec-Santé (Grant #28988).

